# Perfusion via chemicals removes the DNA content of rabbit hearts while keeping the collagen and glycosaminoglycan unaltered

**DOI:** 10.1101/2024.11.22.624857

**Authors:** Banu Seitzhaparova, Sanazar Kadyr, Leya Timur, Baisunkar Mirasbek, Aida Zhakypbekova, Timur Lesbekov, Cevat Erisken

## Abstract

Despite new approaches in the treatment of cardiovascular disease (CVD) such as percutaneous coronary intervention, coronary artery bypass graft, and left ventricular assist devices, which cannot fully compensate for the effectiveness of the original heart, heart transplantation still remains as the most effective solution.

A growing body of literature recognizes the importance of developing a whole heart constructed from living tissues to provide an alternative option for patients suffering from diseases of the cardiovascular system. A potential solution that shows a promise is to generate cell-free, i.e., decellularized, scaffolds using native heart tissue to be later cellularized and transplanted. This study reports the decellularization process and efficiency in an effort to create a whole heart scaffold. The hearts harvested from rabbits were perfused and the final bioartificial scaffolds were characterized for the efficiency of decellularization in terms of DNA content, collagen, and glycosaminoglycan. The compressive biomechanical properties of decellularized and native hearts were also determined and compared. Findings revealed that the DNA content of decellularized hearts was significantly reduced while keeping collagen and GAG content unchanged. Biomechanical properties of the hearth became inferior upon removal of the nuclear material. Decellularized hearts have significant importance in treating CVD as they serve as bioartificial hearts, providing a more clinically relevant model for potential human use.

Future work will focus on the recellularization of the heart using induced pluripotent or embryonic stem cells to test its functionality.

## 1. Introduction

Currently, cardiovascular disease (CVD) is the leading cause of mortality globally, with the risk factors of aging, smoking, overweight, and inadequate exercise in both industrialized and developing countries. These risk factors can lead to mechanical stimuli in the form of pressure overload, and conditions can eventually result with irreversible defects on heart and lead to failure. According to the World Health Organization, over 17 million people die annually from CVDs worldwide[1]. Signs of CVDs are not always obvious and may have similar features to the symptoms of other pathologies unrelated to the heart. Therefore, heart-related dysfunctions remain a significant clinical challenge. Several clinical remedies have been employed to address this challenge, however, with limited success only.[2] These strategies include but are not limited to surgical procedures, angioplasty, valve reconstruction/replacement, and heart transplantation. Hence, there is a need to explore new options over the existing traditional approaches for the prevention and treatment of CVD. Since the decellularized heart retains the spatial array of matrix components, it has various benefits compared to alternative scaffolds. The decellularized extracellular matrix (dECM) plays an important role in the vital activity of the tissue itself and participates in cell communications, division, differentiation, and function. The dECM captures a complex set of proteins such as collagens, GAGs, PGs, and many other ECM compounds that are present in native tissues.[3] In addition, dECM provides signals for the regeneration and restoration of the heart muscles.[4] The dECM retains the structure of the native matrix, and cells can be injected into dECM through hydrogels to restore heart function.

Heart ECM consists of structural and non-structural proteins, which can perform different functions. The main structural proteins are glycoproteins including elastin, fibronectin, laminin, and collagens of types I and III.[5] The amino acid sequence of the polypeptide chains of these proteins makes it possible to form a structure with unique mechanical properties, which have enormous strength and elasticity. Non-structural proteins (proteoglycans) ensure the functioning of dECM as an information center that accumulates and transports signal data for all organ cells. Amongst other things, proteoglycans create a kind of reservoir for cellular factor growth, providing adaptive regenerative capabilities for the heart.

Decellularization is a process aimed at removing cells from the tissues while preserving the extracellular matrix and the three-dimensionality of the organ structure.[6] One of the main goals of successful decellularization is to prevent an immune response upon transplantation by removing nuclear components, damage-associated molecular patterns (DAMPs) and epitopes of the cell membrane, allogeneic or xenogenic DNA.[7]

Existing research recognizes the critical role played by the decellularization approach and its potential outcomes. A chronological improvement in the whole heart decellularization is provided in Table 1. In brief, Ott et al. (2008) is recognized as the first significant attempt to apply perfusion decellularization to rat whole hearts. Then, between 2010 and 2016, the achievements concentrated on trying this protocol on large animals and improving the protocol for increased decellularization efficiency. After 2016 up to date, reduction of decellularization time, increasing perfusion efficiency with the use of new technologies including bioreactors, microfluidics, chips, and automated systems were the major improvements.

**Table 1.** List of decellularization approaches.

| **Year** | **Author** | **Species** | **Improvement** | **Ref.** |
| --- | --- | --- | --- | --- |
| 2008 | Ott | Rat | First whole heart decellularization | [8] |
| 2010 | Wainwright | Pig | Large animal study | [9] |
| 2011 | Weymann | Pig | Use of SDS only at 4% concentration | [10] |
| 2012 | Remlinger | Pig | Increasing trypsin from 0.02% to 0.2%, improving Wainwright's formulation | [11] |
| 2015 | Momtahan | Pig | Increasing decellularization time in Ott's protocol | [12] |
| 2016 | Guyette | Human | Human subject with acceptable level of reduction in DNA*, no loss of mechanical properties and loss of collagens, elastin and GAG after 16days of decellularization | [13] |
| After 2016 |  |  | Reduction of decellularization time, increasing perfusion efficiency with the use of new technologies including bioreactors, microfluidics, chips, and automated systems. |  |
| 2024 | Min | Human | Microfluidic chip device with well-controlled perfusion, acceptable level of DNA removal efficiency*. | [14] |

\* less than 50ng DNA/mg. [14]

Prior reports published on the decellularization of the heart suggest different techniques: chemical, enzymatic, and physical. As described in Table 2, each method has its pros and cons regarding nuclear content removal efficiency, impact on ECM structure integrity, and collagen content.[15] The type of technique to be used is highly dependent upon the type of biological material. Generally, tissues without vasculature are decellularized by immersion and agitation, while vascular tissues require processes like perfusion where cleaning agents are force-penetrated deeper into the tissue. In fact, perfusion decellularization was shown by numerous studies to be able to remove DNA content more efficiently, with an acceptable level of damage to the ultrastructure.[9], [16], [17]

**Table 2.** List of decellularization approaches.

| # | Decellularization type | Solutions | Mechanism of action | Ref. |
| --- | --- | --- | --- | --- |
| 1 | Acidic; Alkaline solutions | Acetic acid; Peracetic acid; Sulfuric acid; Ammonium hydroxide | cell membranes and intracellular organelles are destroyed | [22] |
| 2 | Detergents | Triton X-100; Sodium dodecyl sulfate; SDC; Zwitterionic | Lipids and cell membranes are solubilized | [23] |
| 3 | Antibiotics | Penicillin; Streptomycin; Amphotericin B | Preventing contamination | [24] |
| 4 | Lipases | Gastric lipase; Pancreatic lipase | Solubilization and hydrolysis of lipids | [25] |
| 5 | Proteases | Trypsin; Pepsin; Protease K | Degradation of proteins | [26] |
| 6 | Freeze-thawing | No solutions | Ice crystals formation by freezing cause cell lysing | [27] |

According to Table 2, it is important to note that in most cases, regardless of the method of decellularization, the chemicals used in the process are similar. For example, the non-ionic detergent

Triton X-100 is effective in removing cells and DNA, as it reduces the concentration of glycosaminoglycan (GAGs) and cleaves tissues. Sodium dodecyl sulfate (SDS) is an anionic detergent and is often used in the decellularization of organs to remove cellular components and debris from the tissue. Serine protease-trypsin is used in decellularization to digest cellular proteins hydrolytically and cleaves proteins.[18] Deoxycholic acid decellularizes tissue while preserving structural proteins in an intact position.[19] EDTA is important to prevent blood clotting, and biofilm formation during the decellularization procedure.[18] Peracetic acid is used for disinfection of the tissue while maintaining ECM integrity.[20] Sodium azide is used for bacteriostatic effects to prevent contamination.[21]

Several studies have compared methods of decellularization, studying the elimination of cells and genetic components, and then ensuring the preservation of structural proteins.[23], [28] In this study, chemical and enzymatic methods were utilized to remove DNA content and cell, while keeping intact structural and regulatory proteins by the application of ionic and non-ionic surfactants such as SDS and Triton–X bring solubilization of lipids and cell membranes. Acids and bases with charged properties help with cell membrane solubilization, so peracetic acid and EDTA sterilize and increase the stiffness of ECM. Enzymes such as trypsin help with cell-matrix adhesion damage.[28] Each solution has an impact on ECM in decellularization, nevertheless, it is important to ensure a minimum efficient amount of solution which will not bring side effects for dECM.

From a clinical perspective, using larger animal models in heart decellularization is more relevant. Table 3 lists studies published on heart decellularization of different species. According to Table 3, the time interval varies greatly depending on the size of the heart.

**Table 3.** Review of heart decellularization studies.

| # | Animal model | DNA removal efficiency | Chemicals/Biologicals Used | Duration | Ref. |
| --- | --- | --- | --- | --- | --- |
| 1 | Human heart | n.d. | PBS; NaCl (hypertonic and hypotonic), NaH<sub>2</sub>PO<sub>4</sub> and KH<sub>2</sub>PO<sub>4</sub> in distilled water; SDS; Peracetic acid; NaOH; | 8 days | [31] |
| 2 | The cadaveric rat | 97% | Deionized water; Phosphate-buffered saline; Trypsin; EDTA; Sodium azide; Sodium dodecyl sulfate; Triton-X100; Deoxycholic acid; Peracetic acid; ethanol; | 10 hours | [29] |
| 3 | The cadaveric rat | 96% | Polyethylene glycol; SDS; Triton-X100; | 12 hours | [8] |
| 4 | The cadaveric rat | Protocol I: remove 43%<br>Protocol II: remove 80%<br>Protocol III: remove 99%<br>Protocol IV: remove 99% | Protocol I: PBS with adenosine; SDS; deionized water; Triton-X100;<br><u>Protocol II</u>: Trypsin; EDTA; deionized water; DCA<br><u>Protocol III</u>: Glycerol; NaN<sub>3</sub>; EDTA; DCA; Triton-X100; PBS<br><u>Protocol IV</u>: PBS; SDS; NaN<sub>3</sub>; deionized water; Glycerol; EDTA; DNase I | Protocol I: 6 days<br>Protocol II: 5 hours<br>Protocol III: 12 days and 12 hours<br>Protocol IV: 16 hours | [32] |
| 5 | The porcine heart | n.d. | PBS; deionized water; SDS; Triton-X100; | 7 days | [33] |
| 6 | The porcine heart | 72% | Deionized water; SDS; Triton-X100; | 15 hours | [33] |
| 7 | The cadaveric rat | 99% | PBS with Adenosine; SDS; Triton-X100; Penicillin, Streptomycin and Amphotericin with PBS; | 13 hours 55 min | [34] |
| 8 | The rabbit heart | 87% | EDTA; PBS; SDS; deionized water; ethanol; Sodium azide; | 10 days | [35] |
| 9 | The cadaveric rat | 98.9% | PBS; phenylmethylsulfonyl fluoride; N-ethylmaleimide; benzamidine; iodoacetamide; sodium ascorbate; EDTA; dimethyl sulfoxide; SDS; deionized water; Triton-X100; 2,3-butanedione monoxime; penicillin-streptomycin; amphotericin; | 2 days 21 hours 15 min | [36] |
| 10 | The ovine heart | 88% | SDS; Deoxycholic acid; deionized water; PBC; | 24 hours | [37] |
| 11 | The cadaveric rat | n.d. | PBS; SDS; deionized water; Triton-X100; PBS with penicillin, streptomycin; | - | [38] |
| 12 | The porcine heart | 92% | Trypsin; EDTA; NaN<sub>3</sub>; Triton-X100; deoxycholic acid; peracetic acid; ethanol; | 6 hours 25 min | [9] |
| 13 | The porcine heart | 82% | Trypsin; EDTA; NaN<sub>3</sub>; Triton-X100; deoxycholic acid; peracetic acid; ethanol; | 6 hours 25 min | [10] |
| 14 | The cadaveric rat | n.d. | PBS; SDS; deionized water; Triton-X100; PBS with penicillin, streptomycin, amphotericin; | 13 hours 45 min | [6] |

\*n.d.- not determined

One of the reasons for choosing rabbits as an animal model in this study was that previously rat hearts were decellularized in this group[29] and it is here aimed to scale up the process using a larger animal model. The other reason is the generally recognized similarity between human and rabbit hearts. Marian (2005) compared physiological changes in humans, rabbits, and mice during myocardial tissue damage.[30] Similar heart rates were identified in humans and rabbits. Additionally, from the point of view of the practicality of the study, compared with the human and pig heart, the rabbit heart requires fewer resources and less quantity of reagents for experiments, making it more cost-effective.

The authors of this study previously applied the perfusion decellularization process to rat hearts with successful outcomes. This study, therefore, aimed to decellularize rabbit hearts using perfusion and evaluate the efficiency of the process in terms of its capacity to remove the nuclear component and preserve the structural and non-structural proteins. It is hypothesized that the utilization of the perfusion decellularization method on rabbit hearts would create decellularized whole hearts while keeping their structural properties and non-cellular composition unchanged.

## 2. Materials and Methods

This research study involves two main experimental parts: i) harvesting and decellularization of rabbit hearts, and ii) characterization of decellularized rabbit hearts in terms of DNA content as well as collagen and GAGs. Figure 1 presents the processes for harvesting, decellularization, and characterization of rabbit hearts.

**Figure 1.**
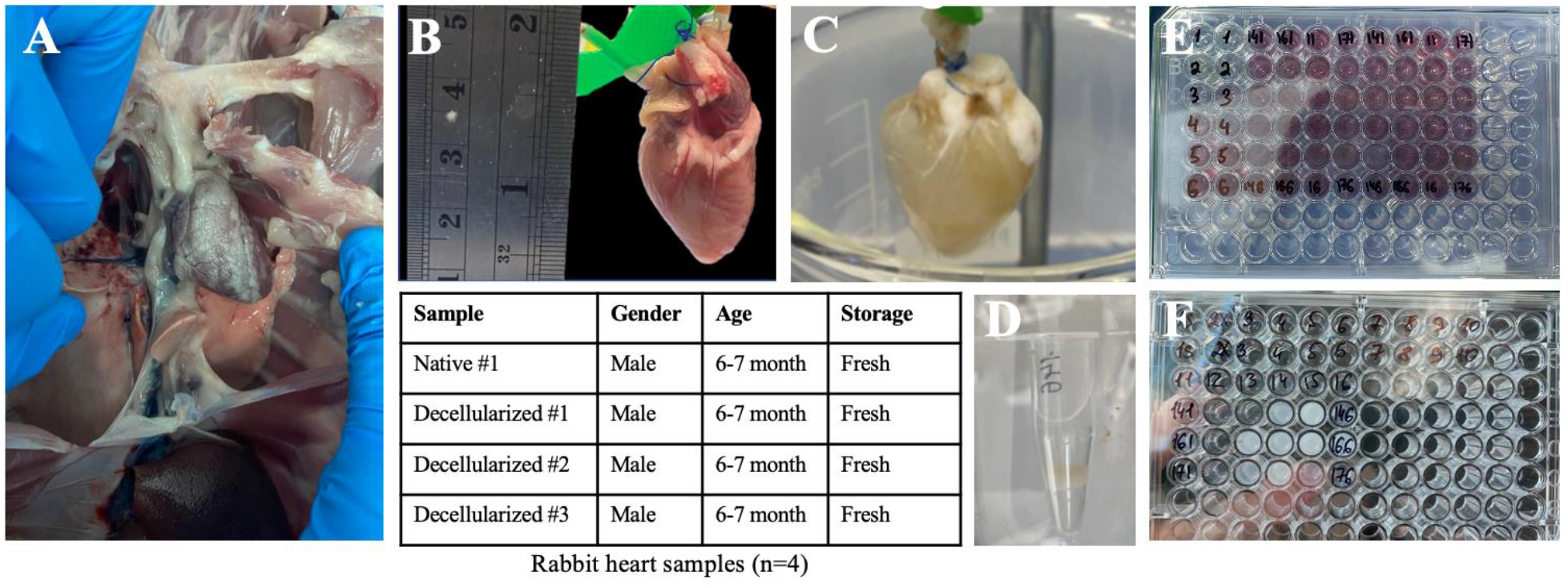
Experimental design and representation of harvesting (A), decellularization (B, C), and characterization (D-F) of rabbit hearts.

### 2.1. Materials

All chemicals were acquired from Sigma Aldrich and Fisher Scientific, and catalog numbers are listed below. Trypsin (Sigma Aldrich, #T4049), Sodium dodecyl sulfate (Sigma Aldrich, #L3771), Sodium azide (Sigma Aldrich, #S2002), Triton-X (Sigma Aldrich, #T9284), Ethylenediamine tetraacetic acid (Sigma Aldrich, #E5134), Peracetic acid (Sigma Aldrich, #433241), ethanol (Sigma Aldrich, #E7023), Phosphate buffered saline (Sigma Aldrich, # P4417), SIRCOL Collagen Standard Type 1 (Sigma Aldrich, #NC9541732), Sodium Formate (Sigma Aldrich, #247596), 1,9-dimethyl methylene blue (Sigma Aldrich, #341088), Sodium acetate trihydrate (Sigma Aldrich, #S7670), L-Cysteine Hydrochloric acid (Sigma Aldrich, #C7477), Papain (Sigma Aldrich, #P3125), PicoGreen double-strand DNA assay (Invitrogen #P7589), chondroitin-6-sulfate (Sigma Aldrich, #C4384).

### 2.2. Harvesting hearts for decellularization

The rabbit hearts were obtained from a local slaughterhouse and harvested along with the aorta and pulmonary artery by the surgical blade (Figure 2A). The perfusion system attached to the hearts through 24G butterfly needles “bioflyject”. To cannulate each heart, the aorta was secured to the end of a cannula using 5.0 Vicryl Plus suture thread. The harvested hearts were used for decellularization and further characterization.

**Figure 2.**
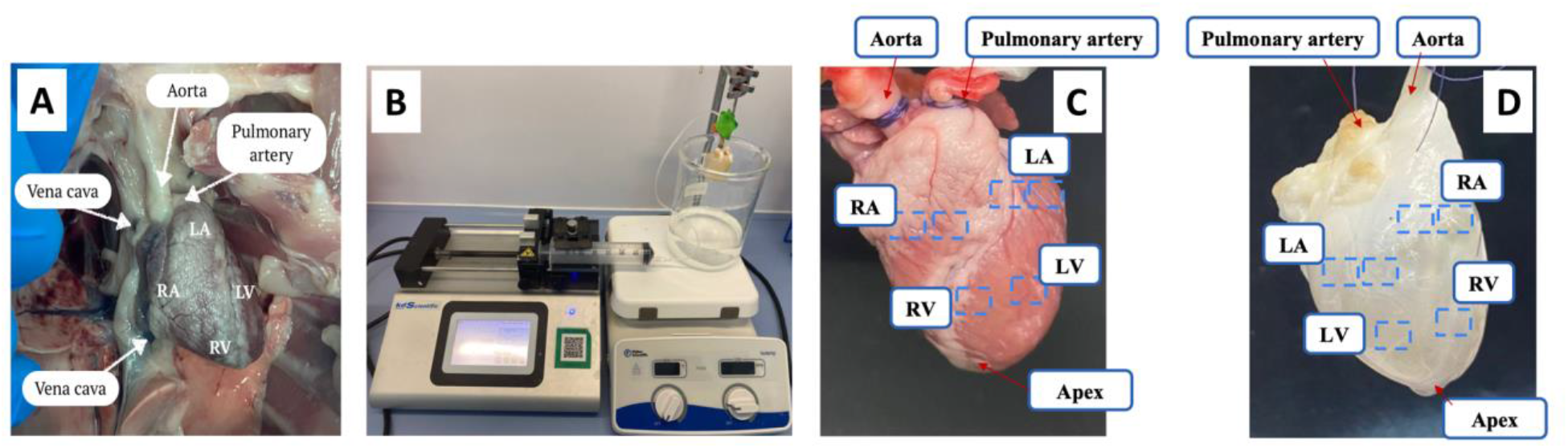
Harvesting and anatomical demonstration of the heart (A), perfusion setup for decellularization (B), and sampling positions for characterization (C, D).

### 2.3. Decellularization procedure

Decellularization of the rabbit hearts was performed according to the modified protocol previously validated on rats by our group[29], with modified durations proportionate to the size of the rabbit heart compared that of the rat. For the perfusion system (Figure 2B), a syringe pump (KDS Legato 210) was used with a configured fluid flow rate parameter of 2.5 ml/min through 60 ml syringes (ID: 26.72 mm). Heart decellularization involved eleven stages, and two cycles were applied to fully remove cellular content.

To compare and ensure effective cell removal, two different perfusions were applied, namely perfusion through the aorta only and bi-ventricular perfusion (Table 4) through both the aorta and pulmonary artery.

**Table 4.** Perfusion procedure for one cycle.

| # | Solution | Aorta Perfusion<sup>a</sup> (mL) | Bi-ventricular perfusion |  |
| --- | --- | --- | --- | --- |
|  |  |  | The aorta<sup>b</sup> (mL) | The pulmonary artery<sup>c</sup> (mL) |
| 1. | Deionized water | 180 | 103 | 77 |
| 2. | Phosphate-buffered saline (PBS) | 360 | 206 | 154 |
| 3. | A mixture of 0.02% trypsin, 0.05% EDTA, and 0.05% Sodium azide, distilled water | 360 | 206 | 154 |
| 4. | A mixture of 1% sodium dodecyl sulfate, and 0.05% Sodium azide, distilled water | 180 | 103 | 77 |
| 5. | A mixture of 3% Triton X-100, 0.05% EDTA, and 0.05% sodium azide, distilled water | 180 | 103 | 77 |
| 6. | A deoxycholic acid, 75% ethanol, distilled water | 180 | 103 | 77 |
| 7. | A mixture of 0.1% peracetic acid, and 90% ethanol, distilled water | 180 | 103 | 77 |
| 8. | PBS | 270 | 154 | 116 |
| 9. | Deionized water | 270 | 154 | 116 |
| 10. | PBS | 270 | 154 | 116 |
| 11. | Deionized water | 270 | 154 | 116 |

a: @2.5 mL/min, b: @1.5 ml/min, c: @1 ml/min

The flow rate of perfusion was 2.5 mL/min for aorta-only perfusion. In bi-ventricular perfusion, perfusion rates through aorta and pulmonary artery were 1.5mL/min and 1mL/min, respectively. Firstly, deionized water of 180 ml was perfused through the aorta to prevent clotting and wash out excess blood. Next, PBS (360 mL) was perfused. After that, the mixture (360 mL) of 0.02% trypsin, 0.05% ethylenediaminetetraacetic acid, and 0.05% sodium azide in deionized water was perfused.

This solution was replaced with a solution (180 ml) of 1% sodium dodecyl sulfate, and 0.05% sodium azide. Next, the mixture (180 ml) of 3% Triton X-100, 0.05% ethylenediaminetetraacetic acid, and 0.05% sodium azide was perfused. The solution (180ml) of deoxycholic acid and 75% ethanol with deionized water was run through the system. This was followed by 180ml of 0.1% peracetic acid, 90% ethanol and distilled water. Next, 270 ml of PBS and 270 ml of deionized water was perfused twice. The bi-ventricular perfusion of rabbit hearts was repeated in the same order of solutions as described in Table 3. The decellularized hearts were stored in PBS at 4°C.

These chemicals were selected and ordered accordingly based on their action on the ECM components. Deionized water and PBS removes blood and blood clot from heart to ease the flow of solutions. The order of trypsin, SDS and Triton-X were selected due to their mechanisms of action. Trypsin-EDTA acts as a protease that cleaves peptide bonds between integral proteins. Trypsin is an ionic surfactant that breaks down the peptides using hydrolysis reaction into amino acid block, and preserves the protein structure.[39] EDTA prevents the integral proteins between cells from binding to one another and disrupting cell adhesion to ECM. Together, trypsin and EDTA is a good combination for decellularization. SDS is an ionic surfactant that works by disrupting non-covalent bonds in the proteins, and so denaturing them, i.e. causing the protein molecules to lose their native conformations and shapes to enable removal of cell nuclei and genetic material.[40] It also removes cytoplasmic proteins from dense tissues. Triton X-100 is a non-ionic surfactant that disrupts the interactions between lipids themselves, and between lipids and proteins. Since it does not disrupt protein-protein interactions, it ensures presence of an intact ECM.[41] Triton X-100 is less effective than SDS but also less destructive due to its non-ionic character. Deoxycholic acid is effective on proteoglycan-associated interstitial network, lyses cells, and solubilizes cellular and membrane components. Peracetic acid removes residual nucleic acids with minimal effect on the ECM composition and structure, and it is a common disinfection agent. The acid was neutralized and removed by perfusing PBS (pH 7.4) and deionized water two times for 30 min. It should be noted that, detergents such as SDS penetrate the tissue, and they must be removed from the remaining ECM structure as quickly as possible. Thus, after using the agents for decellularization, washing the heart tissue with deionized water and PBS is crucial to completely remove detergents from the tissue and neutralize the medium.

### 2.4. Samples preparation

Samples of intact and decellularized hearts were prepared for further examinations (DNA, collagen, GAG assays). A total of 12 pieces of representative tissue specimens were cut from both sides of the heart for analysis (Figure 2C&D). After decellularization, the heart appears as a flat tissue but not as a voluminous organ. From each side, 6 pieces of specimens were collected from different parts of the heart (atria, ventricles, apex) and used for characterization. This is a procedure that was used before.[29], [42]

### 2.5. DNA assay

For a comprehensive analysis of the effectiveness of decellularization, the cellular content of the decellularized heart was analyzed by Picogreen dsDNA quantification kit. The kit was used as recommended by the manufacturer. Samples were quantified using a fluorometric spectrophotometer SkanIt Software 2.4.5 RE for Varioskan Flash with fluorescein excitation (485 nm) and emission (535 nm) wavelengths. A standard curve was created to correlate the DNA content with fluorescence, and the number of cells was determined using the DNA/cell conversion factor. A native intact heart was used as a control group.

### 2.6. Collagen assay

Collagen content of the decellularized hearts was determined using the Sircol Collagen assay kit (Biocolor Ltd, Belfast, N. Ireland). The heart samples were prepared by drying in a vacuum dryer (Buchi Glass Oven B-585) for 22 hours at 40°C. For the colorimetric assay a procedure recommended by the manufacturer was followed. Briefly, Sircol dye reagent was added to samples and standards, and the samples were incubate at 60ºC for 24 hours. Then, upon addition of the digestion buffer (pH 6.0), the samples were centrifuged to pelletize the collagen-dye complex. Afterwards, the liquid supernatant was aspirated and the solid sediment was homogenized with an alkaline solution to separate the combination of collagen and dye. Finally, the absorbance of samples was measured at 540 nm by SkanIt Software 2.4.5 RE for Varioskan Flash. A native intact rabbit heart was used as a control group.

### 2.7. GAG-DMMB assay

To determine the amount of GAGs in the decellularized heart, GAG-DMMB dye-binding assay was utilized, using chondroitin-6-sulfate as a standard. A native intact heart was used for the control group. The intact and decellularized heart samples were prepared by the same procedure as for collagen assay in a vacuum dryer. A digestion buffer (pH 6.0) with Sodium acetate trihydrate, L-Cysteine Hydrochloric Acid, EDTA was added to the tubes containing the samples. Then, 1,9-dimethylmethylene blue dye (pH 1.5) was added to samples and standards. Formic acid was used to reduce the pH level to 1.5. Absorbance of samples was read in the first 5 minutes after adding DMMB dye since the GAG-DMMB complexes precipitate out in 5min and the plate becomes unreadable. The SkanIt Software 2.4.5 RE was used for Varioskan Flash with dual-wavelength at 540 nm (for a positive change) and 595 nm (for a negative change) to improve signal detection.

### 2.8. Biomechanical properties

Biomechanical properties of the native and the decellularized hearts were measured under compression. Specimens were harvested from the hearts using an 8mm diameter punch and placed in between the plates of the instrument (Texture analyzer, TA.XTplusC, Stable Micro System, Hamilton, MA) and compressed at 0.1mm/min. Compressive modulus was calculated using the straight portion of the stress-strain curve. It is the slope of the stress versus strain (change in length divided by normal length) curve.

### 2.9. Statistical analysis

The efficiency of decellularization in terms of DNA, collagen, and GAG contents as well as biomechanical properties was evaluated by comparing it with an intact native heart using a student t-test. A p-value of less than 0.05 was considered a significant difference.

## Results

### 3.1. Visual inspection of decellularized heart

Before examination for cellular content, ECM components, etc., a visual comparison of color change during decellularization was made (Figure 3).

**Figure 3.**
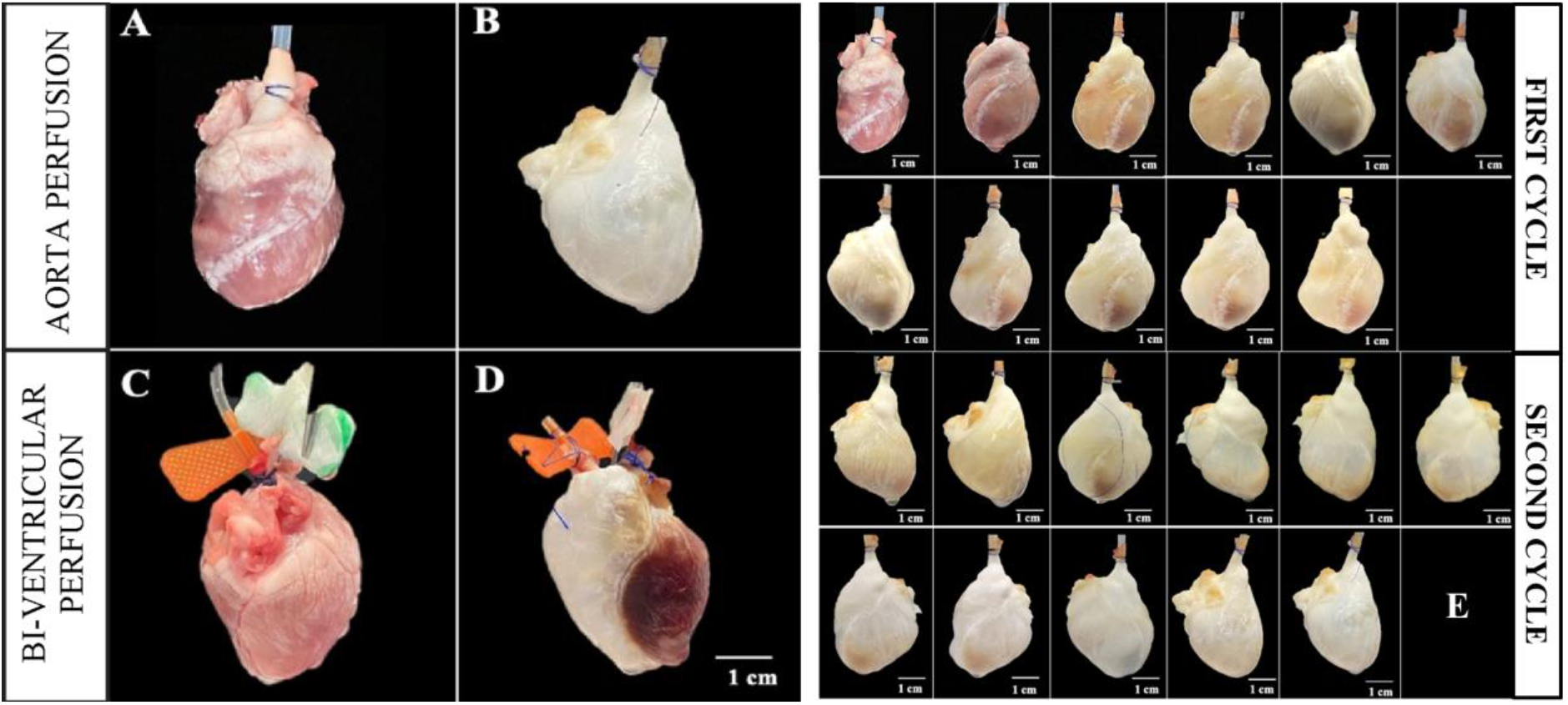
Decellularized heart after aorta (A, B) and bi-ventricular (C, D) perfusion. Stagewise visual quality of aorta perfusion (E) after each step described in Table 3.

Visual comparison after each step of decellularization by aorta perfusion serves as qualitative evidence of the decellularization process as evidenced by a color change from reddish pink in fresh native tissue to translucent white in decellularized tissue (Figure 3A, B, E).

### 3.2. Decellularization by residual nuclear material

A comparison of the results between the native heart and the decellularized heart showed that a significant amount of the DNA content was removed from the hearts upon decellularization (Figure 4A, p<0.05). Approximately 66% reduction in the DNA content was achieved.

**Figure 4.**
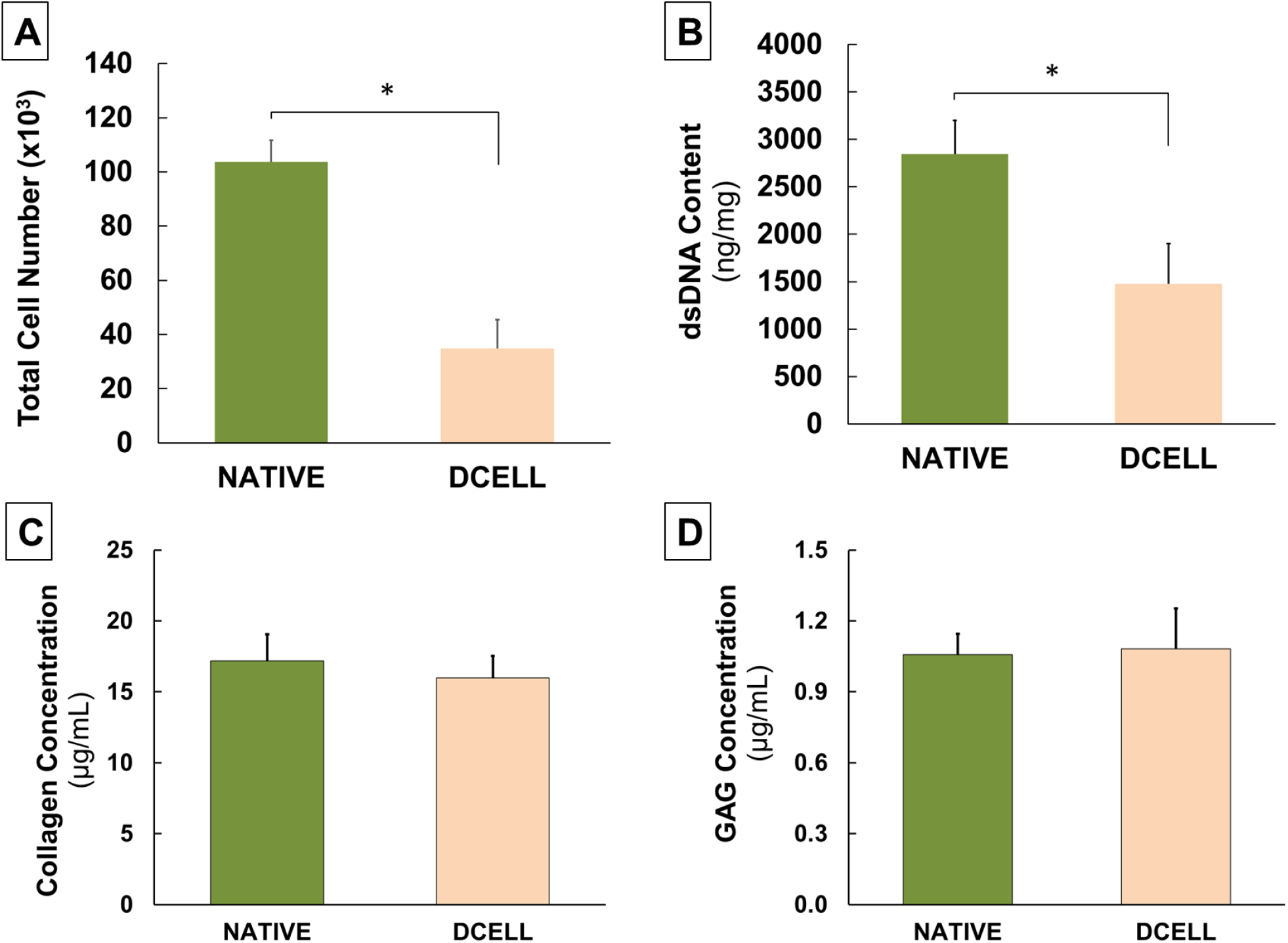
Total cell number (A), dsDNA content (B), collagen content (C) and GAG amount (D) detected in native and decellularized hearts. Error bars represent STD. * indicates difference at p<0.05.

### 3.3. Decellularization assessment by Collagen and GAG assays

Collagen and GAG contents in the decellularized hearts remained unchanged compared to native tissue samples (p>0.05), indicating that the decellularization protocol followed did not remove collagens and GAGs from the heart (Figure 4C and 4D). The protocol was selectively useful for removing the DNA content.

### 3.4. Biomechanical properties

Biomechanical properties of the native and decellularized hearts were determined under compression and reported in the form of stress-strain behavior (Figure 5). Obviously, the removal of nuclear material from the heart led a decrease in the modulus from 47±6.7kPa to 34.2±4.6kPa.

**Figure 5.**
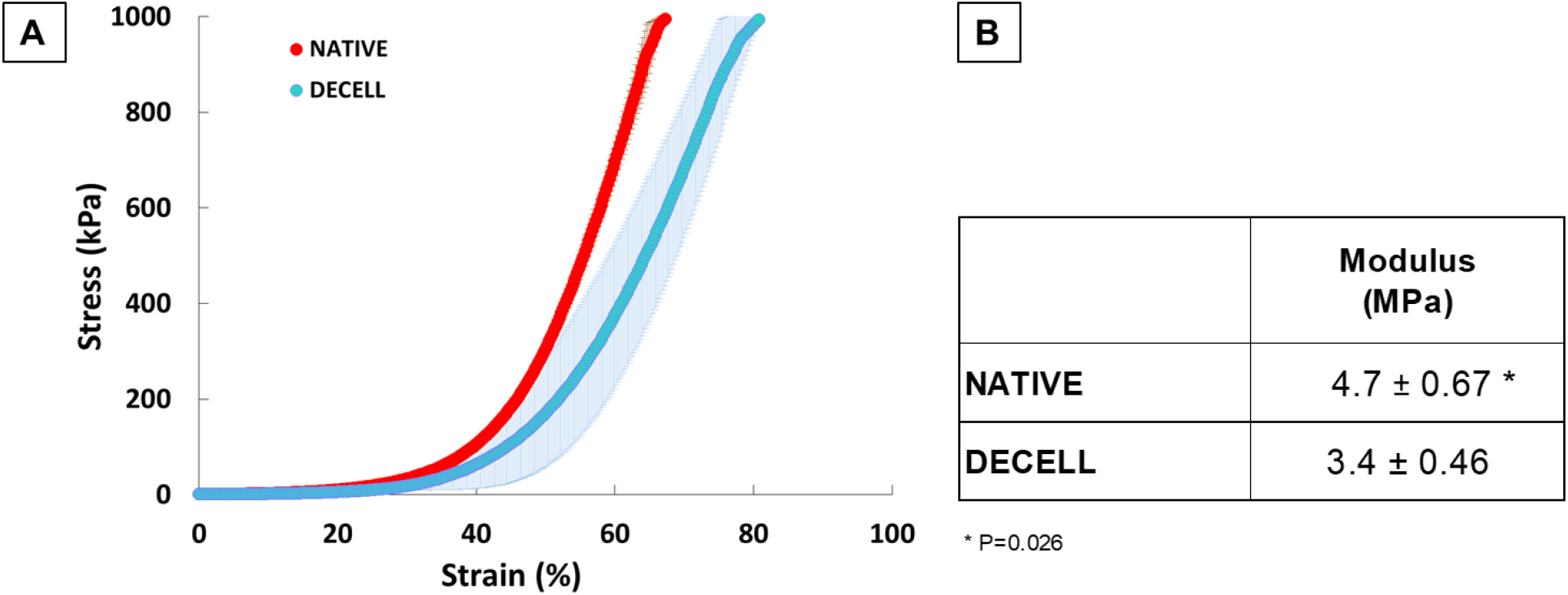
Biomechanical properties of native and decellularized heart. (A) Stress-stress behavior upon compression and (B) Comparison of moduli of native and decellularized heart. Values are in the form of Mean±SD. * represents p<0.05.

## 4. Discussion

The decellularization protocol described in this study was adapted from the cadaveric rat heart decellularization system developed by Ozlu et al., (2009)[29]. The addition in the current study is that two methods of perfusion were applied and compared: perfusion by the aorta and bi-ventricular perfusion. Bi-ventricular and aortic decellularization are two different methods used to create acellular scaffolds. Bi-ventricular decellularization involves the removal of cellular components from the left and right ventricles of the heart, while aortic decellularization involves the removal of cellular components from the aorta, the main artery that carries blood from the heart to the rest of the body. Our experiments using both techniques demonstrated that aorta perfusion is able to clean the heart in two cycle, while in bi-ventricular perfusion the heart was not transparent as a whole. In accordance with heart physiology, blood flow rate in the aorta is much higher than in the pulmonary artery.[43] It is believed that the pressure applied was not sufficiently high to enable the flow of solutions between the two ventricles. Therefore, to save reagents, heart samples, and time, only aortic perfusion was performed.

The primary aim of decellularization is to remove all cellular components while preserving the extracellular matrix; however, complete decellularization is not possible, and the residual cell debris may lead to tissue rejection and calcification.

In this study, a perfusion system was employed to decellularize the entire rabbit heart using various detergents. The effectiveness of detergents in removing cellular material from tissues is attributed to their ability to dissolve the cell membrane and detach proteins from DNA. The effectiveness of the applied rabbit heart decellularization procedure was evaluated by different characterizations. First, a qualitative comparison after each step of decellularization was performed. A color change from reddish pink in fresh native tissue to translucent white in decellularized tissue was observed after the process. The visual observation demonstrated that aorta perfusion resulted in higher decellularization efficiency, and the results were reported based on aorta perfusion. DNA assay using fluorescence spectrophotometry indicated that the DNA content of the heart was significantly reduced as compared to the intact heart. According to previous research on tissue decellularization, the tissue engineering society agreed-upon measurable standards for adequate decellularization. These criteria include less than 50 ng of double-stranded DNA per milligram of extracellular matrix weight; DNA fragment length less than 200 base pairs; no visible nuclear material in tissue sections in staining. [44] In this study, decellularization efficiency was measured in terms of dsDNA content. Our results demonstrated that a 66 ng/mg of DNA level was achieved in the heart samples.

One of the main goals of tissue decellularization is to preserve the maximum integrity of the native structure and components of the extracellular matrix (ECM) to create a scaffold that most accurately mimics the native tissue. As a future work, recellularization and endothelization of the heart are planned, so it is important to preserve the integrity of the ECM. During the endothelization of blood vessels, endothelial cells attach to the basal plate. The main components of the basal plate are type IV collagen, glycosaminoglycans, laminin, fibronectin, elastin, and proteoglycans. Due to the limited time of the study, the contents of only collagen and GAGs were studied. Both collagen and GAG contents in the native and decellularized tissues were found to be similar. In this regard, the decellularization process did not affect the structural integrity of the extracellular matrix or the composition of the tissue.

Biomechanical properties of the whole porcine heart have been previously determined under compression[45], and a significant decrease in modulus was reported after decellularization. Our findings demonstrate that the removal of nuclear material reduced the modulus of the heart despite no change in collagen and GAG contents. Collagen is the major determinant of tissue strength in ligaments and tendons[46], and it also applies to the heart. In this study, we believe that clearing off the cells as well as blood from the heart led to a significant change in the modulus of the heart.

This study certainly has some limitations in the methods applied for the characterization. Therefore, in future studies, for a more accurate assessment of the efficiency of decellularization of the heart, it is recommended to include histological evaluation, quantitative determination of elastin content, the diameter of muscles, and analysis of biomechanical properties. Literature reports suggest that the use of pressure-controlled bioreactors affects the rate of cell leaching from blood vessels. It may also be necessary to extend the continuous decellularization procedure since similar studies on rabbits were conducted continuously for up to 10 days.[35] For recellualirizing the decellularized heart, the dense matrix of cardiac fibers that hinders cellular infiltration should be carefully considered. Investigators in the field researched the ways to promote cell infiltration to find that introducing dynamic flow using a microfluidic chip was very efficient for cellularizing the human whole hearts.[14]

## Conclusion

In this study, the hearts harvested from rabbits were decellularized using a perfusion method. Findings revealed that the DNA content of the decellularized hearts was significantly reduced while keeping collagen and GAG content unchanged. Given the structural similarity between decellularized hearts and native hearts, it is anticipated that the decellularized hearts will provide a more clinically relevant model for potential human use and will have a significant impact in treating cardiovascular diseases.

## Conflict of Interest Statement

Authors declares no conflict of interest.

## Notes

### Competing Interest Statement

The authors have declared no competing interest.

### Summary of Updates

The title was modified to make it more specific for the readers

